# Biobanking of Biological Samples at Room Temperature

**DOI:** 10.1101/2025.04.08.646969

**Authors:** Rajani Kanth Vangala, Saranya Kanukollu, Pratibha M S, Jiny Nair, Anand Babu Vangala, Avinash Shankar, Barsha Upadhyay, Sneha Palpandi, Jeevapriya Nagendran, Pramod N. Nair, Elango E Murugaian

## Abstract

Biospecimens have become an integral part of life science research, pharma and drug development. There is increased demand for biospecimens from patients with assured quality of collection, processing, and storage. Establishing a biobank with high-quality biospecimens is an expensive aspect due to the requirement of costly freezers and liquid nitrogen facilities. Many studies have explored various storage reagents for biological samples. Still, the need of the hour is the one that can preserve biological samples at ambient room temperature and suitable for diagnosis and research. InstaPRESERVE is an eco/user-friendly solution with long-term sustainable alternative for freezer or liquid nitrogen systems for biobanking.

Blood and saliva specimens were collected and stored in InstaPRESERVE at room temperature and conventional storage conditions. The genomics and proteomics analysis of the DNA, RNA, and protein extracted from the -80°C and InstaPRESERVE stored specimens and fresh specimens indicated that the specimens were well-conserved at room temperature compared to frozen specimens.

Innovative technologies and approaches are being evaluated in biobanks for sample integrity, reducing collection, processing, and storage costs, increasing speed and throughput, and reducing energy expenditure. Preserving the biospecimens in InstaPRESERVE at room temperature eliminates expensive storage systems, cold-chain during transportation, and pre-processing requirements. The qualitative and quantitative results of DNA, RNA, and protein ensured stabilized storage for a longer period.

The use of InstaPRESERVE as a specimen storage medium is emerging as an alternative in biobanking for specimen storage at room temperature. The existing and proposed biobanks may consider employing InstaPRESERVE as a reliable resource for biospecimens storage.

## Introduction

Biobanks have become an integral part of life science research and development. The general definition of the biobank is “an organized collection of biospecimens with their associated data for research and clinical applications”(1, 2). Biobanks of human specimens are categorized into disease-oriented and population-based. The disease-oriented banks are hospital-based while population-based are outside the hospitals (3). Biobanks, as specimen resources, are pivotal in basic science, clinical research, and the translation of health knowledge. There is increased demand for large quantities of biospecimens as their associated data are crucial for scientific research and preventive medicine.

Establishing a biobank is expensive due to the requirement of ultra-freezers, data management systems, and liquid nitrogen facilities (4). The complexity of existing biobanks for sample storage and distribution, as well as the risk of man-made errors and disasters, has compromised the integrity of biobank samples and stored data. The maintenance and preventive measures have been established and published by ISBER (5, 6).

The methods for collecting and preserving biological samples include filter papers, Formalin Fixed Paraffin Embedded Tissues (FFPE), polymer matrices, freezing methods, and stabilizing solutions (7-10). Filter paper cards excel in collection but lack downstream processing capability for full nucleic acid recovery (11). FFPE is common in hospitals for preserving tissues for histopathological diagnosis, but poses challenges in proteome investigation due to formalin’s properties (12, 13). Cryopreservation is widely used for long-term cell preservation but risks ice crystal formation and cell damage (14). Lyophilization offers diverse applications but requires expensive equipment and product testing before distribution [10,11]. The polymer matrices are highly versatile and can be adapted to preserve a wide range of biospecimens, including bacteria, plant cells, animal cells, and biomolecules (15). However, the choice of polymer and its application must be carefully tailored to the specific requirements of each type of biospecimen. Tissue-stabilizing solutions are used to collect and transport tissues with varied effects on the biospecimens (16). Formalin is the fixative solution used in clinical pathology. Amber and DMSO are good for preserving DNA, and RNA later for RNA (17, 18). While Amber is eco-friendly, others like formalin, are harmful to the environment and the end users (19-22).

Many studies have explored various storage reagents for biological samples. Still, the need of the hour is the one that can preserve biological samples at room temperature and suitable for diagnosis and research. Tissues were sampled and fixed with six different fixatives for 24 h: formalin, Greenfix, UPM, CyMol, Bouin, and Bouin/Hollande (20). Greenfix is ethanediol and alcohol-based and completely free of formaldehyde, and is the next generation of fixatives. UPM, which is a combination of ethanol, methanol, isopropyl alcohol, and formalin; and CyMol, which is a combination of ethanol, methanol, and isopropyl alcohol. The results indicated that UPM and CyMol may be regarded as potential substitutes for formaldehyde with the possibility of technical improvement and standardization of protocols.

InstaPRESERVE^™^ is an eco-friendly solution to preserve biospecimens at room temperature. This biodegradable and user-friendly product is suitable for all types of biospecimens for the collection, transport, and clinical and research analysis (23). The specimens are well preserved and ideal for histopathology, genomics, and proteomics studies. It is proposed as an appropriate medium for biobanking, wherein expensive infrastructure like deep freezers and liquid nitrogen are not required. This also eliminates the use of biohazard chemicals in specimen collection and storage. The application could be extended to various fields, including medical diagnostics, forensics, agriculture, veterinary medicine, and life science research. This study evaluated the application of InstaPRESERVE^™^ in biobank specimen storage and genomics and proteomics studies.

## Materials and Methods

### Biobank specimen collection

Biospecimens, blood and saliva samples from 20 volunteers (10 aliquots of 200 µl per sample) were collected and stored in InstaPRESERVE^™^ at room temperature and the control aliquots were stored at -80°C. The storage containers were sealed properly to avoid any leakage. These specimens were stored for over 36 months.

### Genomics study

The genomics evaluation of the samples was conducted at 6 months,18 months, and 36 months. The control and IP samples of blood and saliva were subjected to DNA and RNA extraction. For every extraction, one fresh sample (blood/saliva) was included. The DNA from whole blood was isolated using the lysis buffer (24) and TRIzol (Invitrogen) method. For salivary DNA isolation, the Polgarova and coworkers’ method was followed (25). Total RNA was extracted with the Hybrid-RTM kit (GeneAll Biotechnology Co., Ltd, South Korea). The qualitative and quantitative analysis of DNA and RNA was assessed using EPOCH spectrophotometer. The DNA and RNA samples were visualized by 0.8% agarose gel electrophoresis. The DNA and RNA samples were further evaluated by Cyp2d6 PCR using specific primers (26). The amplicons were visualized by 1% agarose gel electrophoresis.

The DNA samples were evaluated for their suitability in next-generation sequencing. The cancer hotspot panel, Ion AmpliSeq Cancer Hotspot Panel v2 (Ion Torrent Gene Studio S5, Thermo Fisher, USA), was used. The primary analysis of sequencing read lengths corresponds directly to the quality of the DNA and the sequencing fragments generated. The read length data were analyzed using a Student’s t-test to determine statistical significance.

### Protein extraction and SDS PAGE

The proteins were isolated by TRIzol method from the biospecimens estimated by Bradford method (27). These proteins were stored at -80°C, and further analyzed by SDS PAGE.

## Results

### Biobanking

The samples stored in InstaPRESERVE™ for more than 36 months did not exhibit any deterioration, discoloration, or precipitation (Fig. 1). There was no contamination in any of the samples stored. The control blood, saliva, and tissue samples stored at -80°C were also intact. The results have indicated that blood and saliva specimens stored with InstaPRESERVE™ at room temperature are suitable for biobanking.

**Figure 1.**
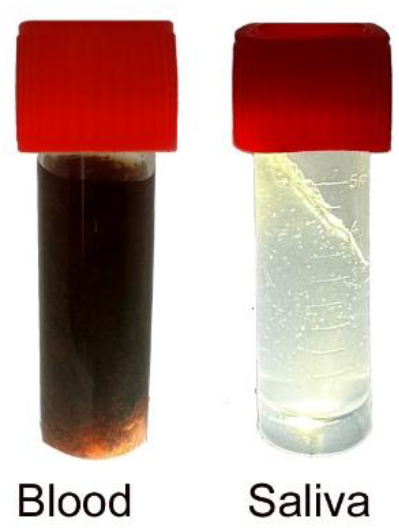
Biobank biospecimens (blood and saliva) stored in InstaPRESERVE™.

### Genomics analysis

The quantitative analysis of the DNA and RNA extracted from specimens stored at -80°C and InstaPRESERVE™ is presented in Table 1 and Figure 2. When compared to fresh specimens, the yield obtained was lower for the samples stored at -80°C and InstaPRESERVE™. While the percentage yield of -80°C specimens decreased at 18 months and 36 months, the yield was stabilized for specimens stored with InstaPRESERVE™. This indicated that the specimens were well preserved at room temperature. The gel electrophoresis analysis exhibited that good-quality DNA and RNA were extracted from all specimens (Figures 3 and 4). The Cyp2D6 PCR results further confirmed their suitability for molecular analysis (Figure 5).

**Table 1:** Quantitative values of DNA, RNA, and protein isolated from the biobank specimens.

| S. No | Description | Fresh specimen | 6 months |  | 18 months |  | 36 months |  |
| --- | --- | --- | --- | --- | --- | --- | --- | --- |
| | | | $-80^{\circ}\text{C}$ | IP | $-80^{\circ}\text{C}$ | IP | $-80^{\circ}\text{C}$ | IP |
| **DNA (ng/$\mu\text{l}$)** | Blood | 480.48 | 337.25 | 200.34 | 326.21 | 223.4 | 212.23 | 194.96 |
|  | Saliva | 201.44 | 188.47 | 125.2 | 123.08 | 100.68 | 130.34 | 91.65 |
| **RNA (ng/$\mu\text{l}$)** | Blood | 208.76 | 134.44 | 145.61 | 118.91 | 102.8 | 108 | 86.19 |
|  | Saliva | 53.01 | 44.22 | 35.63 | 42 | 39.94 | 39.51 | 38.24 |
| **Protein (ug/mL)** | Blood | 7.68 | 7.03 | 7.14 | 5.46 | 5.04 | 3.94 | 3.52 |
|  | Saliva | 3.02 | 2.71 | 2.83 | 2.29 | 2.18 | 1.8 | 1.38 |

**Figure 2:**
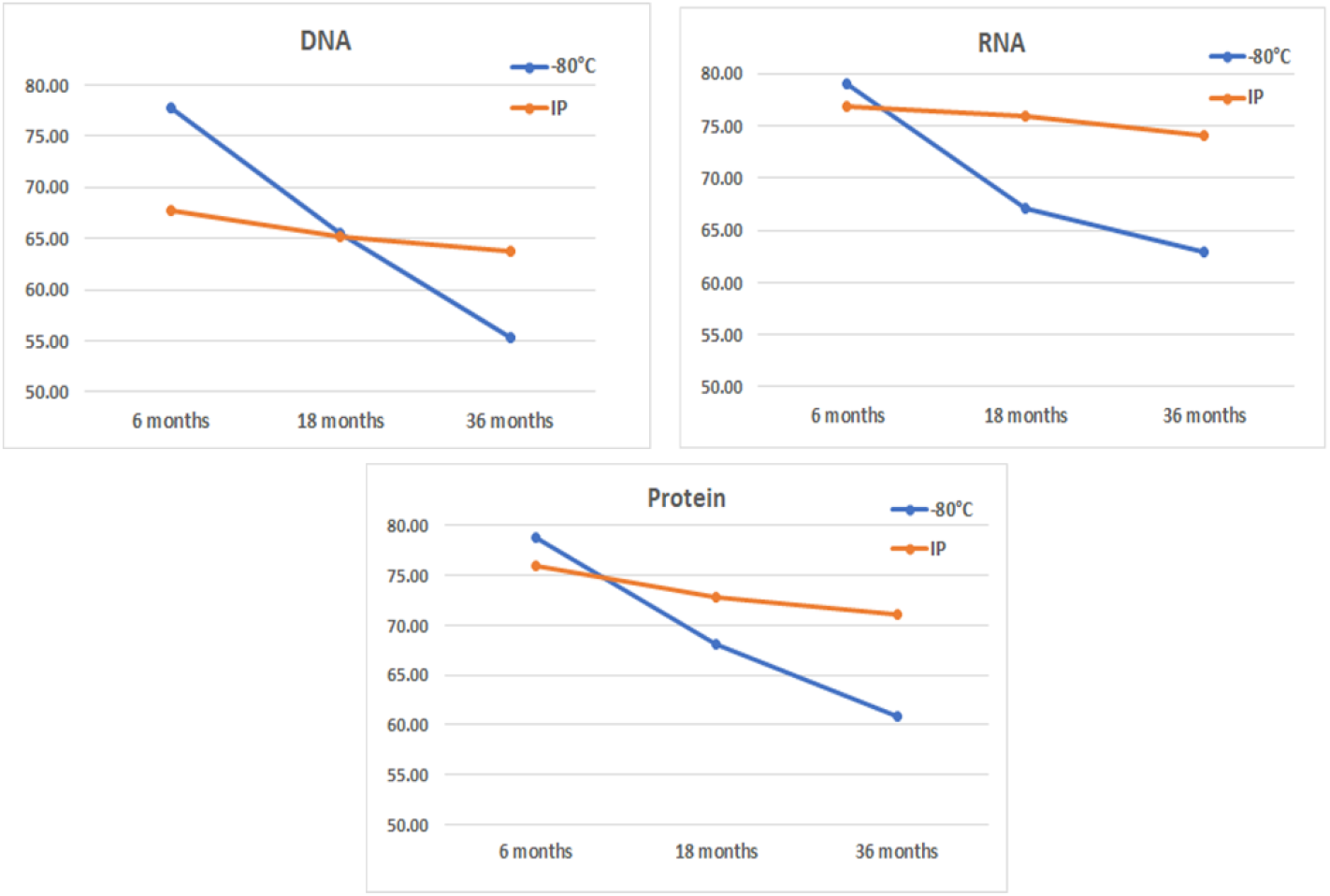
The percentage yield of DNA, RNA, and protein. The amount of DNA, RNA, and protein obtained from the biobank specimens at three different time points and the storage conditions were compared with those of the fresh specimens.

**Figure 3.**
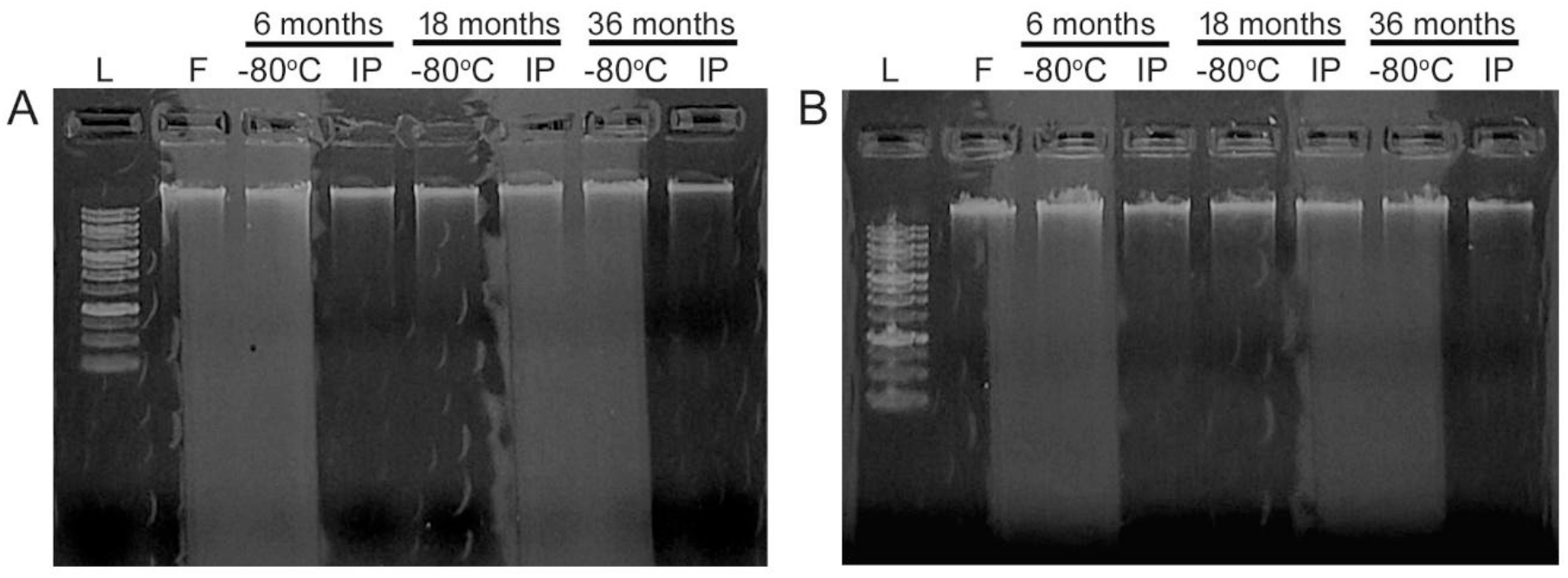
Stability Study of DNA. The DNA isolated from the biospecimens was analyzed by agarose gel electrophoresis. **(A)** Blood, **(B)** Saliva. Lane 1: 100 bp Ladder, Lane 2: ‘F’ Fresh sample. Lanes 3 & 4: Samples stored for 6 months at -80°C and InstaPRESERVE™, Lanes 5 & 6: Samples stored for 18 months at -80°C and InstaPRESERVE™, Lanes 7 & 8: Samples stored for 36 months at -80°C and InstaPRESERVE™.

**Figure 4.**
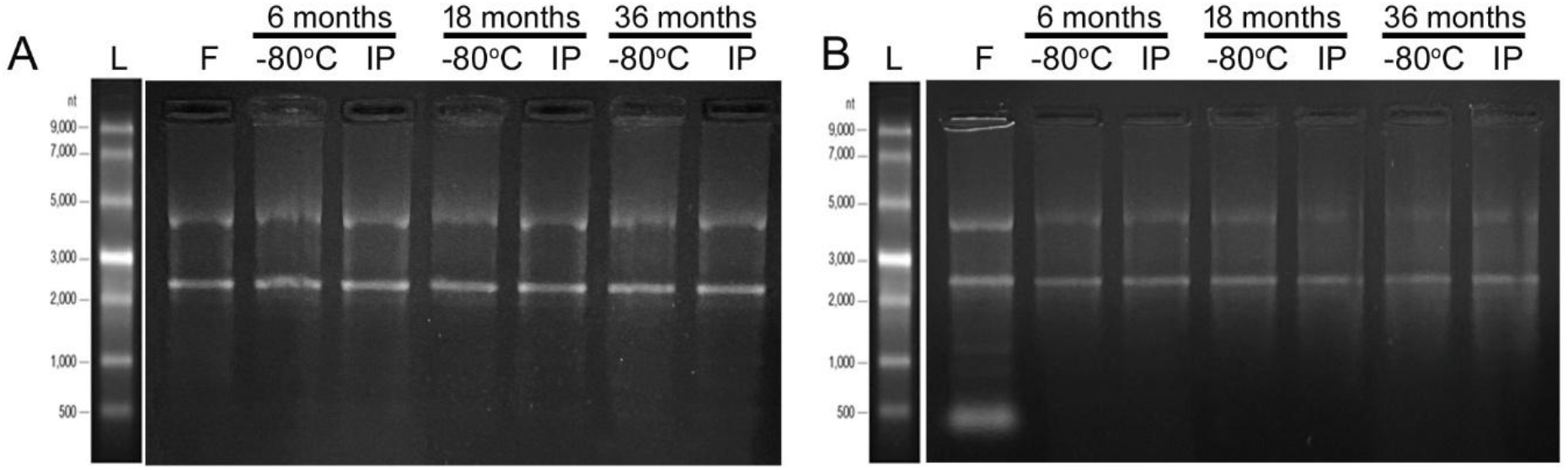
Stability Study of RNA. The RNA isolated from the biospecimens was analyzed by agarose gel electrophoresis. **(A)** Blood **(B)** Saliva. Lane 1: RNA Ladder, Lane 2: ‘F’ Fresh sample. Lanes 3 & 4: Samples stored for 6 months at -80°C and InstaPRESERVE™, Lanes 5 & 6: Samples stored for 18 months at -80°C and InstaPRESERVE™, Lanes 7 & 8: Samples stored for 36 months at -80°C and InstaPRESERVE™.

**Figure 5.**
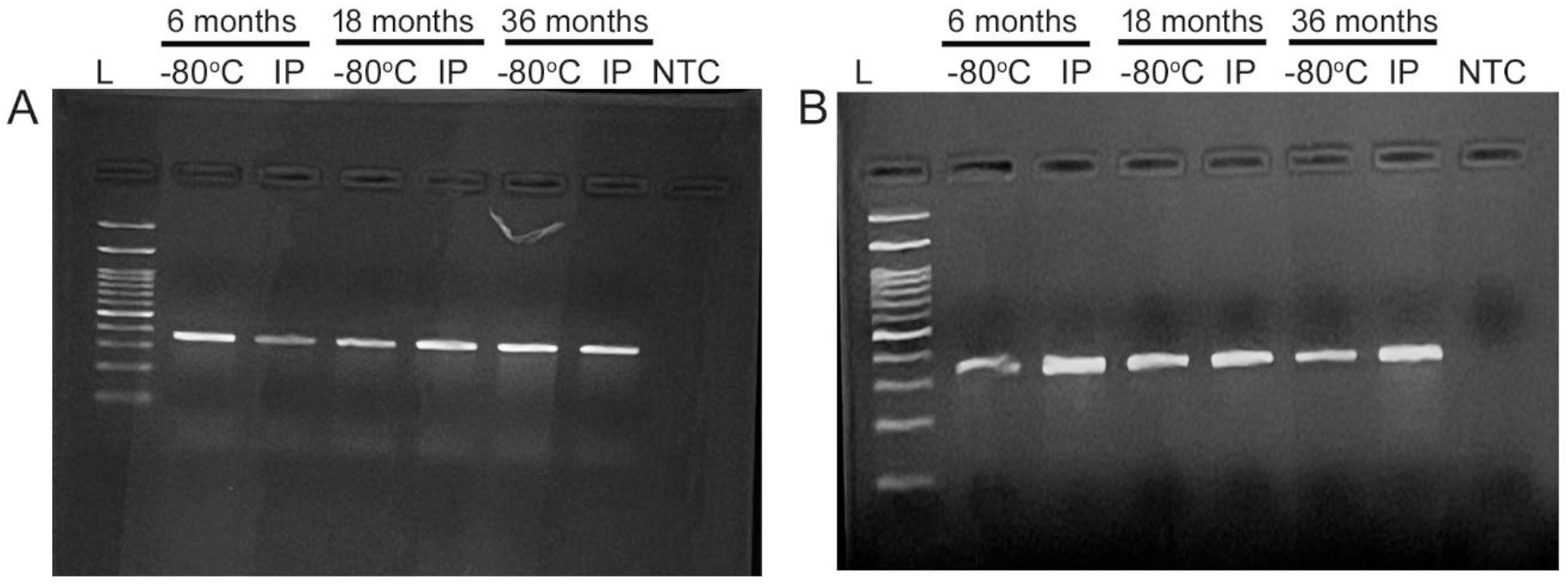
Cyp2D6 PCR analysis of the biospecimens’ DNA. The DNA isolated from the biospecimens was amplified with Cyp2D6 PCR primers. **(A)** Blood **(B)** Saliva. Lane 1: DNA Ladder, Lanes 2 & 3: Samples stored for 6 months at -80°C and InstaPRESERVE™, Lanes 4 & 5: Samples stored for 18 months at -80°C and InstaPRESERVE, Lanes 7 & 8: Samples stored for 36 months at -80°C and InstaPRESERVE™.

The results obtained from Ion AmpliSeq Cancer Hotspot Panel v2 confirmed that the DNA samples were suitable for next-generation sequencing (Table 2, Figure 6). The t-test analysis of the read length data from blood and saliva revealed that instaPRESERVE stored specimens’ DNA quality did not significantly differ from fresh specimens and frozen specimens (blood p value 0.37936; saliva p value 0.49966)

**Table 2:**
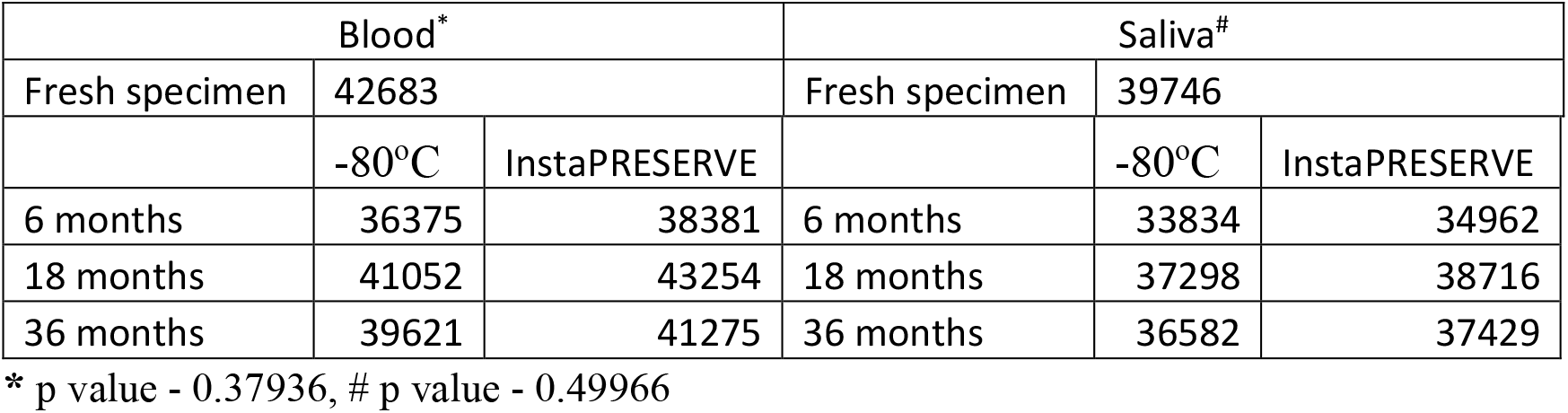
Sequencing reads obtained from the Ion AmpliSeq Cancer Hotspot Panel v2 amplification of the DNA samples.

| Blood* |  |  | Saliva# |  |  |
| --- | --- | --- | --- | --- | --- |
| Fresh specimen | 42683 |  | Fresh specimen | 39746 |  |
|  | -80°C | InstaPRESERVE |  | -80°C | InstaPRESERVE |
| 6 months | 36375 | 38381 | 6 months | 33834 | 34962 |
| 18 months | 41052 | 43254 | 18 months | 37298 | 38716 |
| 36 months | 39621 | 41275 | 36 months | 36582 | 37429 |

\* p value - 0.37936, # p value - 0.49966

**Figure 6.**
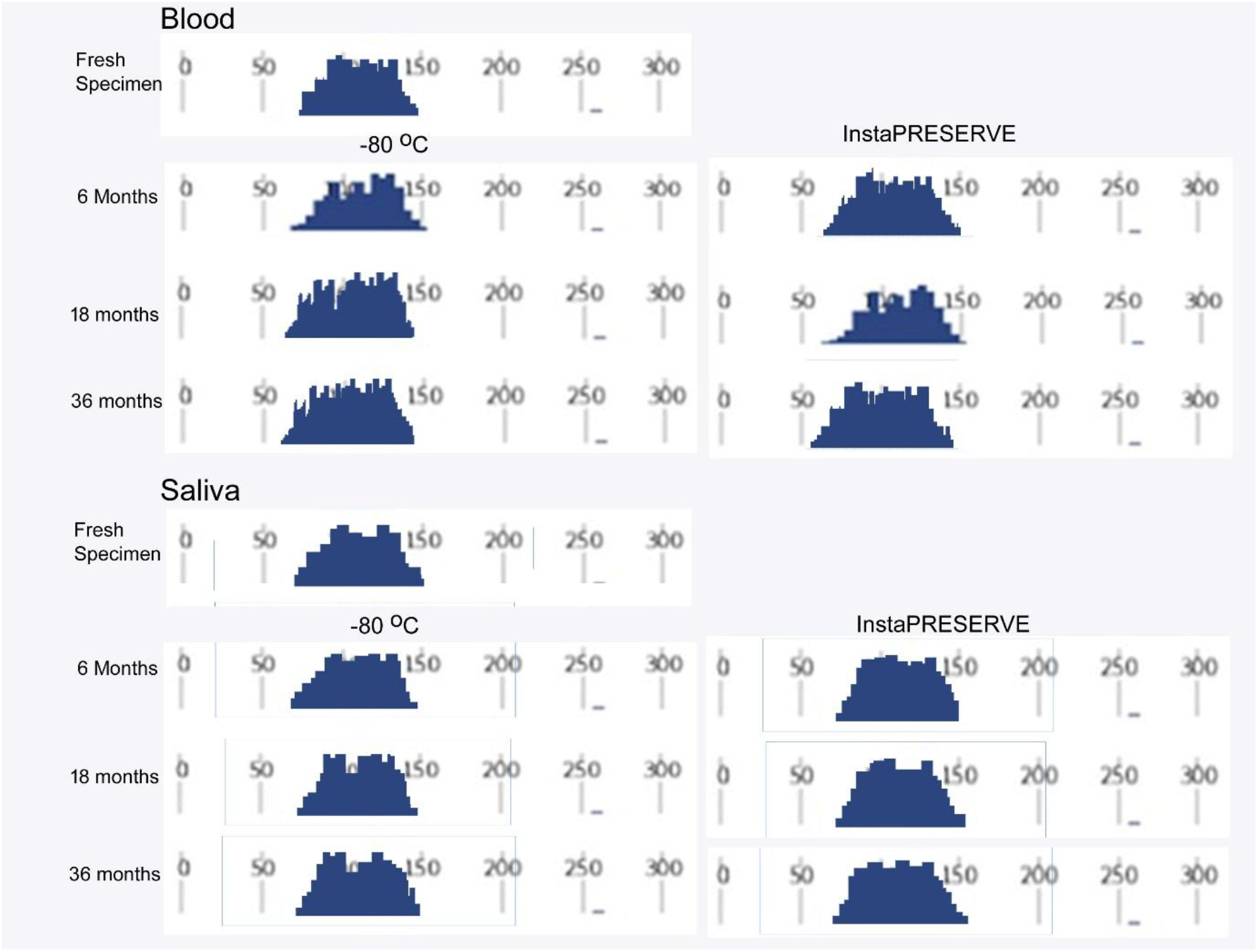
Ion AmpliSeq Cancer Hotspot Panel v2 reads analysis of the biospecimens’ DNA. The DNA isolated from the biospecimens was subjected to Ion AmpliSeq Cancer Hotspot Panel v2 analysis.

### Protein analysis

The amount of total protein extracted from the biobank specimens is presented in Table 1. As discussed above, the percentage of protein yield obtained from InstaPRESERVE™ stored specimens was lower but stable, while -80°C stored specimens yielded a decrease with the storage period. The electrophoresis results also showed that the quality was not compromised for both storage conditions (Figure 7).

**Figure 7.**
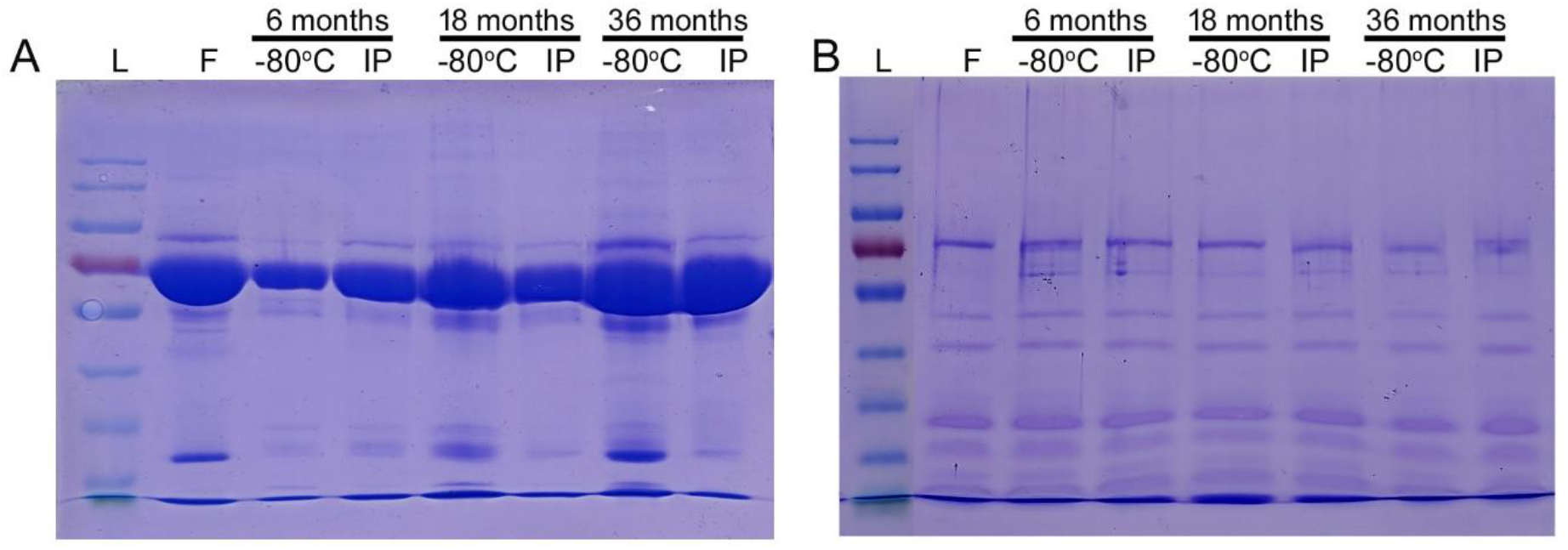
Protein stability Study. The total protein isolated from the biospecimens was analyzed by polyacrylamide gel electrophoresis. **(A)** Blood **(B)** Saliva. Lane 1: RNA Ladder, Lane 2: ‘F’ Fresh sample. Lanes 3 & 4: Samples stored for 6 months at -80°C and InstaPRESERVE™, Lanes 5 & 6: Samples stored for 18 months at -80°C and InstaPRESERVE™, Samples stored for 36 months at -80°C and InstaPRESERVE™.

## Discussion

Biobanks are indispensable biomedical research facilities for clinical, drug discovery, preclinical, and translational studies (28). A successful biobank is highly dependent on the support of health professionals, infrastructure, and financial support (29). The biospecimen quality has been a neglected area in biobanks. To address this, the National Cancer Institute, USA, has established a biospecimen research database (30). The biospecimen quality is evaluated by histopathology for tumor, integrity of DNA and RNA for genomics, and protein for proteomics studies (31, 32). The mode of biospecimen collection, preprocessing, storage, and transport is the key factor, depending on the type of specimens collected (33). One of the key factors in tumor banking is the ischemia time: warm ischemia time from blood vessel ligation to surgical excision time, and the cold ischemia time from excision to freezing. These factors influence the gene expression profiles (34).

Innovative technologies and approaches are being evaluated in biobanks for sample integrity, reducing collection, processing, and storage costs, increasing speed and throughput, and regulating energy expenditure (35). The existing methods for sample collection, transport, and storage rely on formalin-fixed tissues, frozen specimens of blood, saliva, urine, and body fluids. This involves toxic and carcinogenic chemicals like formalin and expensive instruments like freezers and liquid nitrogen containers. The cryopreservation and cryobiology are known to be far from adequate (36). Due to limitations, the available alternative methods have restricted use in biobanks (31). InstaPRESERVE™ is an alcohol-based, biodegradable, and user-friendly solution evaluated for its application in biobanking. Preserving biospecimens in InstaPRESERVE™ at room temperature eliminates expensive storage systems, cold chain during transportation, and pre-processing requirements. The specimens are fixed during collection; hence, the genomics and proteomics profiles are well-conserved, as represented by our results. In tissues, especially in tumors, the cold ischemic condition is eliminated. The yield percentage results of DNA, RNA, and protein emphasize that stabilized storage for a longer period is ensured.

## Conclusion

The evolving field of biobanking is revolutionized with innovative approaches and technologies for sample collection, storage, integrity, and reduced energy expenditure. The existing methods employ toxic chemicals such as formaldehyde, high-energy-consuming freezers, and liquid nitrogen containers. Earlier reported alternative techniques and fixatives have not reached a comprehensive solution. The innovation of InstaPRESERVE™ as a specimen storage medium with fixation property is emerging as an alternative in biobanking for specimen storage at room temperature. The results obtained are conclusive of InstaPRESERVE™ as an apt resource for cancer research. The limitations are that this has been validated with blood, saliva, and tissues of different organs for genomics, proteomics, and histopathology. This product has to be studied for other biochemical and metabolic parameters and specimens, like urine and body fluids. The existing and proposed biobanks may consider employing InstaPRESERVE™ as a reliable resource for biospecimens storage.

## Declaration of competing interest

The authors declare that there is no conflict of interest.

## Acknowledgement

Part of this study was funded by BIRAC, Department of Biotechnology, Government of India

## Notes

### Competing Interest Statement

The authors have declared no competing interest.

